# OSCA/TMEM63 are an Evolutionarily Conserved Family of Mechanically Activated Ion Channels

**DOI:** 10.1101/408732

**Authors:** Swetha E. Murthy, Adrienne E. Dubin, Tess Whitwam, Sebastian Jojoa-Cruz, Stuart M. Cahalan, Seyed Ali Reza Mosavi, Andrew B. Ward, Ardem Patapoutian

## Abstract

Mechanotransduction, the conversion of mechanical cues into biochemical signals, is crucial for many biological processes in plants and animals^1,2^. In mammals, some mechanosensory processes such as touch sensation and vascular development are mediated by the PIEZO family of mechanically activated (MA) ion channels^3-5^. In plants, the impact of gravity or soil properties on root development, wind on stem growth, and turgor pressure on plant-cell size and shape are proposed to involve activation of MA ion channels^6,7^. Homologues of the bacterial MA channel MscS (MSLs) exist in plants, and MSL8 is shown to be involved in pollen hydration^8^; however, the identity of the MA channels required for most mechanotransduction processes in plants have remained elusive^9^. Here, we identify various members of the 15 OSCA proteins from *Arabidopsis thaliana* (previously reported as hyperosmolarity sensors^10,11^) as MA ion channels. Purification and reconstitution of OSCA1.2 in liposomes induced stretch-activated currents, suggesting that OSCAs are inherently mechanosensitive, pore-forming ion channels. This conclusion is confirmed by a high-resolution electron microscopy structure of OSCA1.2 described in a companion paper^12^. Beyond plants, we present evidence that fruit fly, mouse, and human TMEM63 family of proteins, homologues of OSCAs, induce MA currents when expressed in naïve cells. Our results suggest that OSCA/TMEM63 proteins are the largest family of MA ion channels identified, and are conserved across eukaryotes. We anticipate that further characterization of OSCA isoforms which have diverse biophysical properties, will help gain substantial insight on the molecular mechanism of MA ion channel gating and permeation. OSCA1.1 mutant plants have impaired leaf and root growth under stress, potentially linking this ion channel to a mechanosensory role^11^. We expect future studies to uncover novel roles of OSCA/TMEM63 channels in mechanosensory processes across plants and animals.

## Main

In *Arabidopsis thaliana,* hyperosmolarity-evoked intracellular calcium increase is dependent on the genes OSCA1.1 and OSCA1.2^10,11^. However, the activation mechanism for these proteins and whether they encode a pore-forming ion channel remains unknown. We synthesized human codon optimized versions of OSCA1.1 (At4g04340) and OSCA1.2 (At4g22120) cDNA in pIRES2-mCherry vector, heterologously expressed them in mechanically-insensitive PIEZO1-knockout HEK293T cells (HEK-P1KO)^13,14^, and electrophysiologically characterized hyperosmolarity-activated currents. In contrast to published reports^10,11^, we find that hyperosmolarity-evoked whole-cell currents recorded from OSCA1.1- or OSCA1.2-expressing cells were only modestly larger than baseline currents (**Extended Data Fig.1**).

We next explored the possibility that OSCA1.1 and OSCA1.2 are mechanosensitive, and that the modest hyperosmolarity-induced currents might be due to osmotic shock causing cell shrinking, and affecting membrane tension^15^. In cells, MA currents are commonly induced by two direct methods: 1) cell-membrane indentation with a glass probe induces macroscopic MA currents in the whole-cell patch clamp mode; 2) cell-membrane stretch induces single-channel or macroscopic MA currents when pressure is applied to a recording pipette in the cell-attached (or excised) patch clamp mode. Surprisingly, MA whole-cell currents recorded from cells transfected with OSCA1.1 or OSCA1.2 were 10-100-fold larger (10-fold for OSCA1.1 and 100-fold for OSCA1.2, **Fig. 1a** vs. **Extended Data Fig. 1**) than the hyperosmolarity-activated currents, and were comparable to those recorded from cells transfected with mouse PIEZO1, a well-characterized mechanosensitive ion channel (**Fig. 1b**). OSCA1.1 and OSCA1.2 whole-cell MA currents had an apparent activation threshold of 8.6 ± 0.9 µm and 6.3 ± 0.7 µm, and inactivated (channel closure in continued presence of stimulus) with a time constant of 10.0 ±1.3 ms and 10.4 ± 1.7 ms, respectively (**Fig. 1b** and **Extended Data Table. 1**). Similarly, robust macroscopic stretch-activated currents were recorded from cells transfected with OSCA1.1 or OSCA1.2 but not from mock-transfected cells (**Fig. 1c, d**). Stretch-activated currents from OSCA1.1 and OSCA1.2 were reversible and inactivated with a time constant of 24 ± 3.4 ms and 24.6 ± 4.8 ms, respectively (**Fig. 1d**). The pressure required for half-maximal activation (P_50_) of OSCA1.1 and OSCA1.2 was 58.5 ± 3.7 mmHg and 54.5 ± 2.2 mmHg, respectively (**Fig. 1e**). These values are higher than mouse PIEZO1 which has a threshold of 24 ± 3.6 mmHg^3,16^ (**Fig. 1e**), demonstrating that at least in HEK-P1KO cells these proteins evoke high-threshold MA currents. These results suggest that OSCA1.1 and OSCA1.2 are involved in mechanotransduction.

**Figure 1.**
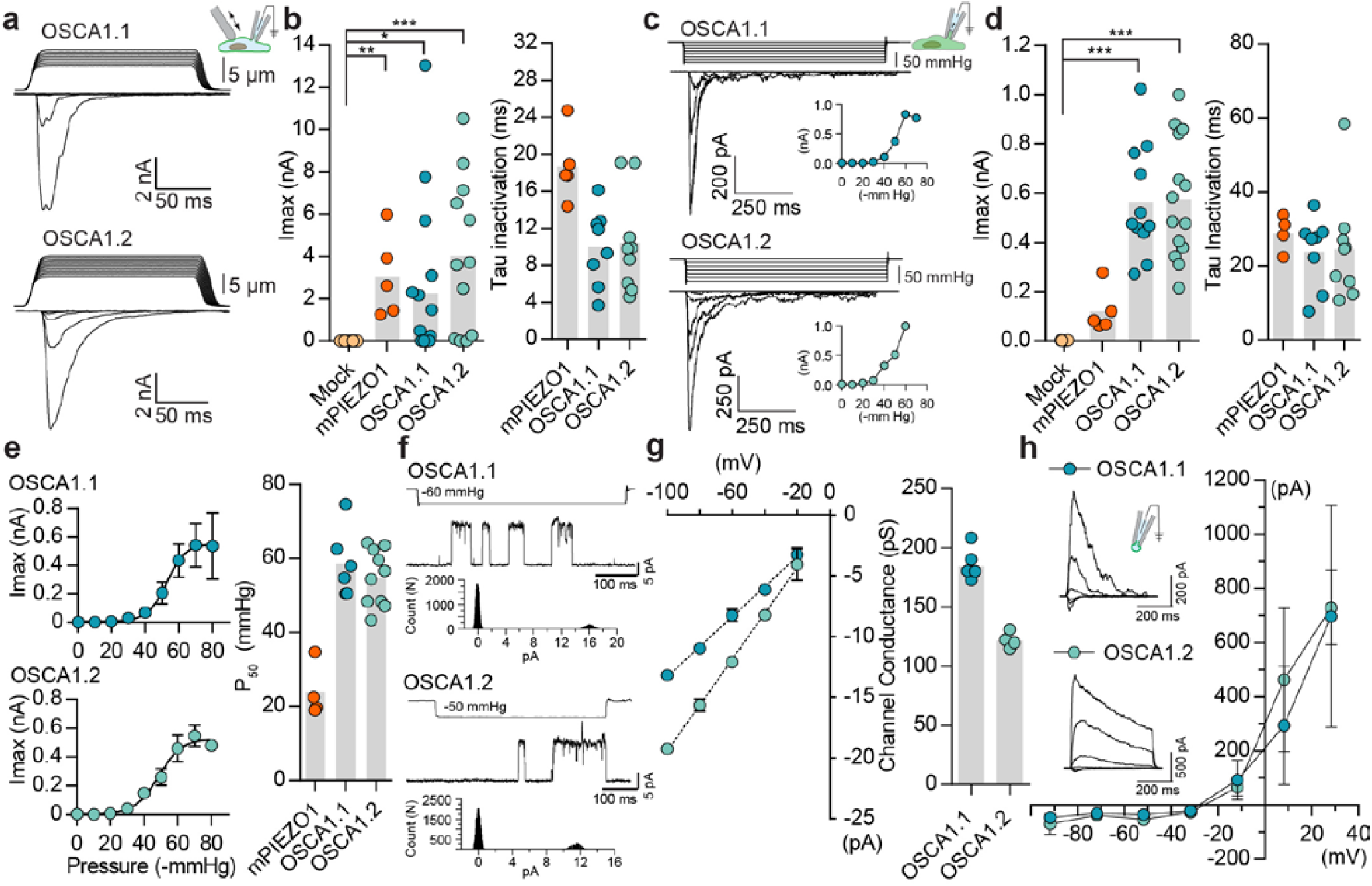
OSCA1.1 and 1.2 induce MA currents in HEK-P1KO cells. **a,** Representative traces of MA whole-cell currents (−80 mV) from OSCA1.1- and OSCA1.2-expressing cells. The corresponding probe displacement trace is illustrated above the current trace. **b,** Left, indentation-induced maximal currents recorded, before the patch is lost, from HEK-P1KO cells expressing mock plasmid (N=10), mouse PIEZO1 (N=5), OSCA1.1 (N=16, 9 gave responses) or OSCA1.2 (N=12, 10 gave responses). Right, Inactivation time constant (ms) for individual cells across mPIEZO1 (N=5), OSCA1.1 (N=8), and OSCA1.2 (N=9) (\**P*=0.013, \*\**P*=0.005, \*\*\**P*<0.0001, Kruskal-Wallis test). **c,** Representative traces of stretch-activated macroscopic currents (−80 mV) from OSCA1.1- and OSCA1.2-expressing cells. The corresponding pressure stimulus trace is illustrated above the current trace. Inset represents pressure-response curve for the representative cell. **d,** Left, Maximal currents recorded from HEK-P1KO cells expressing mock plasmid (N=7), mPIEZO1 (N=5), OSCA1.1 (N=11), or OSCA1.2 (N=14). Right, Inactivation time constant (ms) for individual cells across mPIEZO1 (N=5), OSCA1.1 (N=8), and OSCA1.2 (N=9) (OSCA1.1: \*\*\**P*=0.0005, OSCA1.2: \*\*\**P*=0.0001, Kruskal-Wallis test). **e,** Average pressure-response curves (fit with a Boltzman equation) for stretch-activated currents from mPIEZO1-(N=4), OSCA1.1-(N=6), or OSCA1.2-(N=10) expressing cells. Bar graph on the right represents P50 values for individual cells across the two isoforms. **f,** Representative single-channel currents (−80 mV) recorded in response to negative pipette pressure as indicated. Amplitude histogram for the trace is illustrated below. Channel openings are upward deflections. **g**, Left, average I-V relationship of stretch-activated single-channels from OSCA1.1- and OSCA1.2-transfected cells. Right, Mean channel conductance for individual cells across OSCA1.1 (N=5) and OSCA1.2 (N=4). **h,** Average I-V of stretch-activated currents recorded from outside-out patches excised from cells expressing OSCA1.1 (Erev:-24.5±3.3, N=4) or OSCA1.2 (ERev:-25.7±0.7, N=5). Inset: representative current traces.

Under physiological ionic conditions, OSCA1.1- and OSCA1.2-dependent stretch-activated currents had single-channel conductance of 184 ± 4 pS and 122 ± 3 pS, respectively (**Fig. 1f, g**). We note that these values are larger than what was previously reported^11^. Unlike our measurements, the single-channel conductance described in Yuan et al were measured in the absence of a stimulus, and might not be strictly OSCA-dependent. Interestingly, stretch-activated single-channel current traces from either protein revealed a single sub-conductance state, which was half the amplitude of the full open state (**Extended Data Fig. 2**). Sub-conductance states are a hallmark of many ion channels and are indicative of concerted gating of multiple pores from the same channel, or of changes within a single pore^17-19^. The presence of a single intermediate state in OSCA1.1 and OSCA1.2 could be suggestive of two cooperative subunits. Indeed, this suggestion is confirmed by the cryo-EM structure of OSCA1.2, which reveals a pore within each subunit of a dimeric channel^12^. Characterization of the stretch-activated currents in asymmetrical NaCl solution revealed that OSCA1.1 and OSCA1.2 evoked non-selective cation currents with some chloride permeability (OSCA1.1: P_Cl_/P_Na_= 0.21 ± 0.06; OSCA1.2: P_Cl_/P_Na_= 0.17 ± 0.01) (**Fig. 1h**). In addition, gadolinium, a generic MA cation channel blocker, inhibited OSCA1.1- and OSCA1.2-induced stretch-activated currents (**Extended Data Fig. 3**). Together, these results demonstrate that OSCA1.1 and OSCA1.2 induce MA non-selective cation currents.

In *Arabidopsis thaliana*, OSCA1.1 and OSCA1.2 belong to a family of genes that include 15 isoforms across 4 clades^10,11^ (**Fig. 2a**). OSCA1.1 and OSCA1.2 share 85% sequence identity, while other isoforms within the same clade are more divergent (50-70%). Isoforms within clade 2 have about 30% identity to Clade 1 isoforms, and Clade 3 and 4 share the least homology with Clade 1 (**Fig. 2a, 2c**, **Extended Data Table. 2**). We investigated whether mechanosensitivity might be conserved across this family. We selected one gene from each clade, and tested whether they can induce MA currents in a heterologous expression system (**Fig. 2**). In the whole-cell patch clamp mode, mechanical indentation of cells expressing OSCA1.8, OSCA2.3, OSCA3.1, or OSCA4.1 isoforms did not elicit MA currents (**Extended Data Fig. 4**). Remarkably, however, distinct stretch-activated currents were recorded from cell expressing OSCA1.8, OSCA2.3, or OSCA3.1, but not OSCA4.1 (**Fig. 2b**, **Extended Data Table. 1**). Although the lack of OSCA4.1-induced MA currents could suggest that this isoform is functionally distinct from the other isoforms, we cannot rule out that OSCA4.1 might be incorrectly folded or not trafficked to the membrane in HEK-P1KO. The three mechanosensitive isoforms had disparate gating kinetics, with different inactivation time constants (**Fig. 2c**). While the pressure of half-maximal activation was comparable among all the mechanosensitive OSCA clones (**Fig. 2d**), the most striking feature was the diversity in their single-channel conductance (**Fig. 2e, f**). OSCA1.8 and OSCA3.1 channel conductance was approximately four- and six-fold smaller than OSCA1.1, respectively. Furthermore, single-channel amplitude of OSCA2.3 stretch-activated currents were in the sub-picoampere range and unresolvable, which would explain the relatively smaller maximal current responses (**Fig. 2c**). These results provide evidence that OSCAs are a family of MA ion channels with unique biophysical properties. Intriguingly, the isoforms have differential responses to the two types of mechanical stimulation. It is noteworthy that OSCA1.1 and OSCA1.2 which were the only isoforms activated by membrane indentation, had a high threshold (8 µm) compared to mouse PIEZO1 (4-5 µm)^13^ (**Extended Data Table. 2**). Perhaps other OSCA isoforms have even a higher threshold, technically limiting us from recording reliable MA currents, since HEK-P1KO cells break at 10-12 µm of indentation. Alternatively, mechanical indentation and membrane stretch deliver biophysically-distinct forces to the membrane and the different isoforms might be tuned to the unique type of force applied.

**Figure 2.**
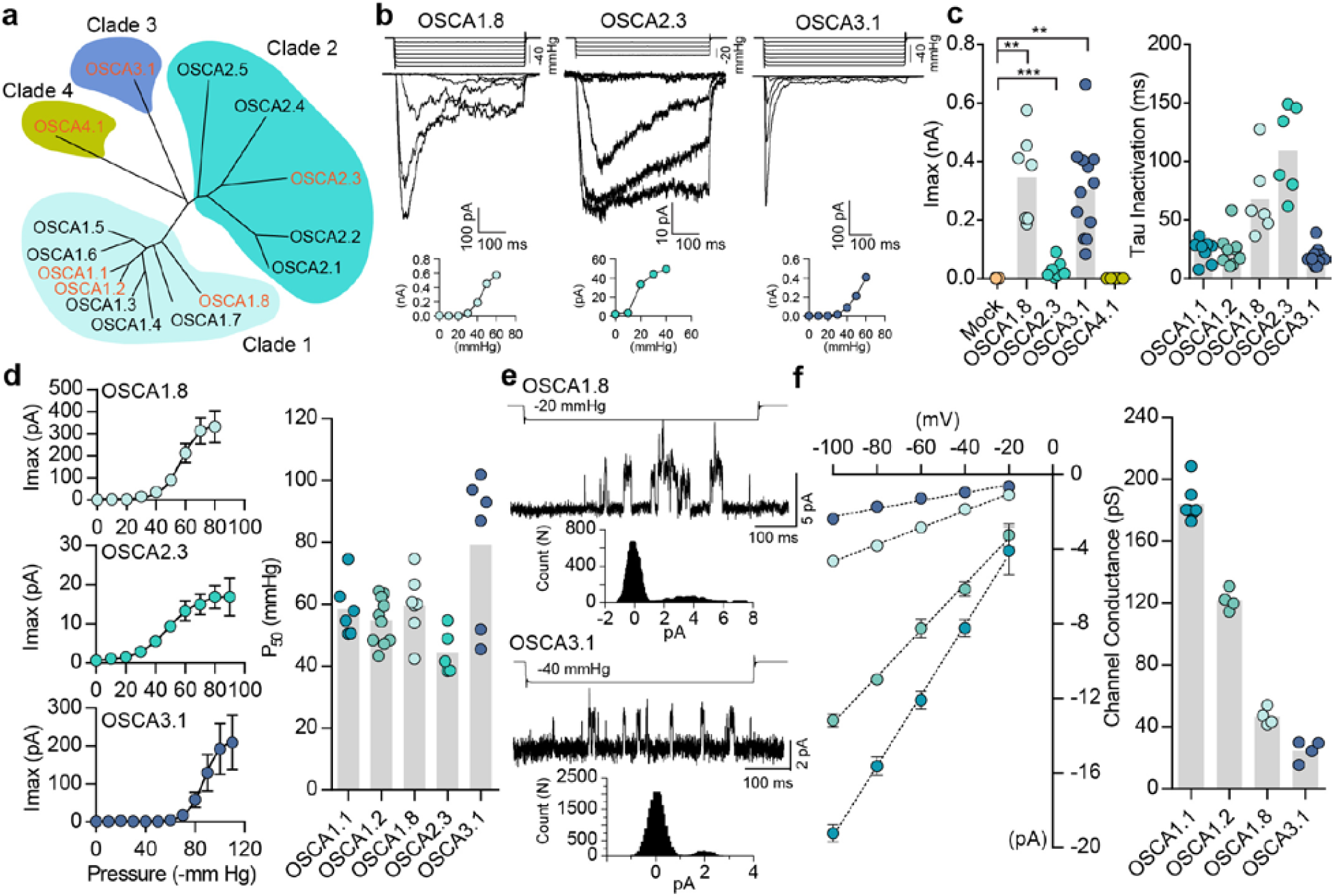
Distinct subclasses of OSCA family members induce MA currents in HEK-P1KO cells. **a,** Phylogenetic tree describing sequence relationship between the 15 OSCA isoforms. Protein sequences were aligned using MegAlign Pro and tree was generated using DrawTree. Isoforms in orange were selected for characterization of mechanically-induced biophysical properties of the channel. **b,** Representative traces of stretch-activated macroscopic currents recorded from cells expressing OSCA1.8, OSCA2.3, or OSCA3.1. The corresponding stimulus trace is illustrated above the current traces. Pressure-response curve for the representative cell is illustrated below the traces. **c,** Left, maximal current recorded from individual cells expressing mock plasmid (N=7), OSCA1.8 (N=7), OSCA2.3 (N=7), OSCA3.1 (N=12), or OSCA4.1 (N=6). Right, inactivation time constant for individual cells from OSCA1.1 (N=8), OSCA1.2 (N=9), OSCA1.8 (N=6), OSCA2.3 (N=6), and OSCA3.1 (N=11) (OSCA1.8: \*\**P*=0.004, OSCA3.1: \*\**P*=0.003, OSCA2.3:\*\*\**P*=0.0006, Kruskal-Wallis and Mann-Whitney tests). **d,** Average pressure-response curves fit with Boltzmann equation for OSCA1.8, OSCA 2.3 and OSCA3.1. Individual P_50_ values for cells from each isoform are plotted on the right (OSCA1.1 (N=6), OSCA1.2 (N=10), OSCA1.8 (N=7), OSCA2.3 (N=5), and OSCA3.1 (N=6)). **e**, Representative stretch-activated single-channel currents recorded at −80 mV from cells expressing OSCA1.8 or OSCA3.1. Channel openings are upward deflections. The stimulus trace for the current is illustrated above. Amplitude histogram for the trace is depicted below. **f**, Average single-channel I-V curves and slope conductance for the indicated isoforms. OSCA1.1: N= 5, OSCA1.2: N= 4, OSCA1.2: N=4, and OSCA3.1: N=4. OSCA1.1 and OSCA1.2 data from Fig.1 is replotted in this figure for comparison.

The results described so far indicate that OSCAs evoke stretch-activated currents in a heterologous expression system, meeting at least one of the criteria for MA channels. Indeed, these results by themselves only allow us to conclude that they are a component of an ion channel; it is formally possible that they require endogenously expressed components within HEK-P1KO cells to behave as a functional mechanosensitive ion channel. To address this possibility, we purified OSCA1.2 fused to a GFP tag, and reconstituted the protein in liposomes to determine whether stretch-activated currents can be recorded from excised proteoliposome patches (**Fig. 3a**)^2^. Purified OSCA1.2-GFP (117 kDa) on a denaturing gel appears as a single protein band, indicating the absence of other associated subunits (**Extended Data Fig. 5**). Before characterizing OSCA1.2, we first tested whether we could successfully record stretch-activated currents from liposomes reconstituted with the bacterial mechanosensitive ion channel *E. coli* MscL^20^. We observed MscL-like stretch-induced channel activity in seven out of eight excised patches (**Extended Data Fig. 5** and **Fig. 3b-d**). The stretch-activated single-channel currents had a conductance of 3340 ± 180 pS (N=6), which is in accordance with previous reports^21^ (**Fig. 3c**). Remarkably, applying negative pipette-pressure in the range of 0 to 50 mmHg to patches excised from liposomes reconstituted with OSCA1.2-GFP protein also induced robust macroscopic currents (**Fig. 3e, g**). Applying negative pressure (as high 100 to 120 mmHg) to patches excised from empty liposomes did not change the baseline current. At the single-channel level, the stretch-activated currents exhibited sub-conductance states, were voltage-dependent, and had a conductance of 346 ± 13 pS in 200 mM KCl solution, which matched the single-channel conductance measured from OSCA1.2-expressing cells (304 ± 9 pS) under the same ionic concentration (**Extended Data Fig. 5** and **Fig. 3f, h**). These results demonstrate that OSCA1.2, like other bona-fide MA ion channels such as MscL, TREK and TRAAK, and PIEZO1, is directly gated by changes in membrane tension, and requires no additional cellular components for activation^20,22-24^. This represents conclusive evidence that OSCA1.2 encodes a pore-forming subunit and is inherently mechanosensitive^12^.

**Figure 3.**
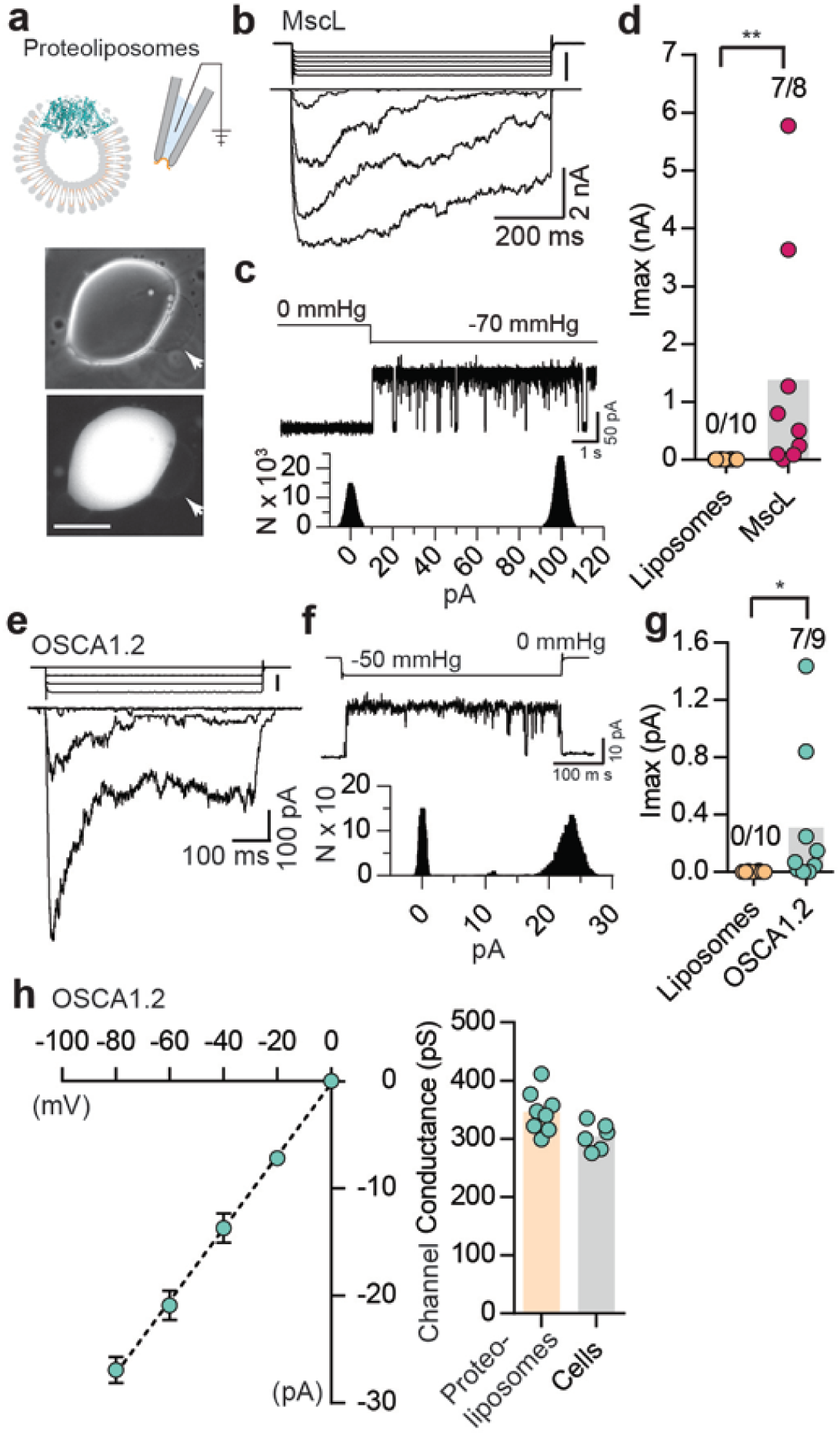
Reconstituted OSCA1.2 in liposomes form MA ion channels. **a,** Illustration to depict that patches were pulled from liposomes reconstituted with purified MscL or OSCA1.2 proteins. Channels in patch were activated by negative pipette pressure. Brightfield (top) and GFP (bottom) images of proteoliposomes reconstitutes with OSCA1.2-GFP protein. Patches were pulled from unilamellar vesicles (indicated by arrows). Scale bar: 50 µm. **b,** Representative stretch-activated currents from unilamellar liposomes reconstituted with EcMscL. The corresponding negative pipette pressure-stimulus is illustrated above the current traces. **c,** Representative single-channel trace recorded in response to −70 mmHg pressure. Channel openings are upward deflections. Currents were filtered at 10kHz. Amplitude histogram of the full-trace is depicted below. **d,** Maximal stretch-activated currents recorded from empty liposomes or EcMscL reconstituted liposomes. Fractions indicate attempts that resulted in currents/total number of attempts (\*\**P*=0.003, Mann-Whitney test). **e**-**g,** Liposomes reconstituted with OSCA1.2-GFP protein. Layout is same as b-d. Single-channel currents in **f** were filtered at 2 kHz. Also note the difference in channel amplitude relative to EcMscL. **g**, Maximal stretch-activated currents recorded from empty liposomes or OSCA1.2-GFP (\**P*=0.04, Mann-Whitney test). **h,** Left, average I-V relationship of stretch-activated single-channels in liposomes reconstituted with OSCA1.2-GFP. Right, single-channel conductance in 200mM KCl solution from individual proteoliposome patches (N=8) or cells expressing OSCA1.2 (N=6). In **c** and **f** N represents count or number of events.

We next examined if orthologues of OSCAs are also mechanosensitive. Phylogenetic analysis and previous studies have identified the TMEM63 family of proteins as the closest homologues of OSCAs^10,11,25^ (**Fig. 4a**). To determine whether mechanosensitivity was conserved across different species, we selected isoforms in fruit fly, mouse, and human, and tested their ability to induce MA currents in HEK-P1KO cells. In the whole-cell patch clamp mode, mechanical stimulation of cells expressing DmTMEM63, MmTMEM63A, MmTMEM63B, MmTMEM63C, or HsTMEM63A did not elicit MA currents (**Extended Data Fig. 4**). However, these clones (with the exception of MmTMEM63C) induced stretch-activated currents when expressed in naïve cells (**Fig. 4b,c**). The phylogenetic tree illustrates that mammalian TMEM63C is divergent from TMEM63A and TMEM63B (**Fig. 4a**), which may explain its lack of mechanosensitivity. Alternatively, it is possible that this protein is incorrectly folded and not trafficked to the membrane, or that TMEM63C is not sufficient by itself to induce MA currents.

**Figure 4.**
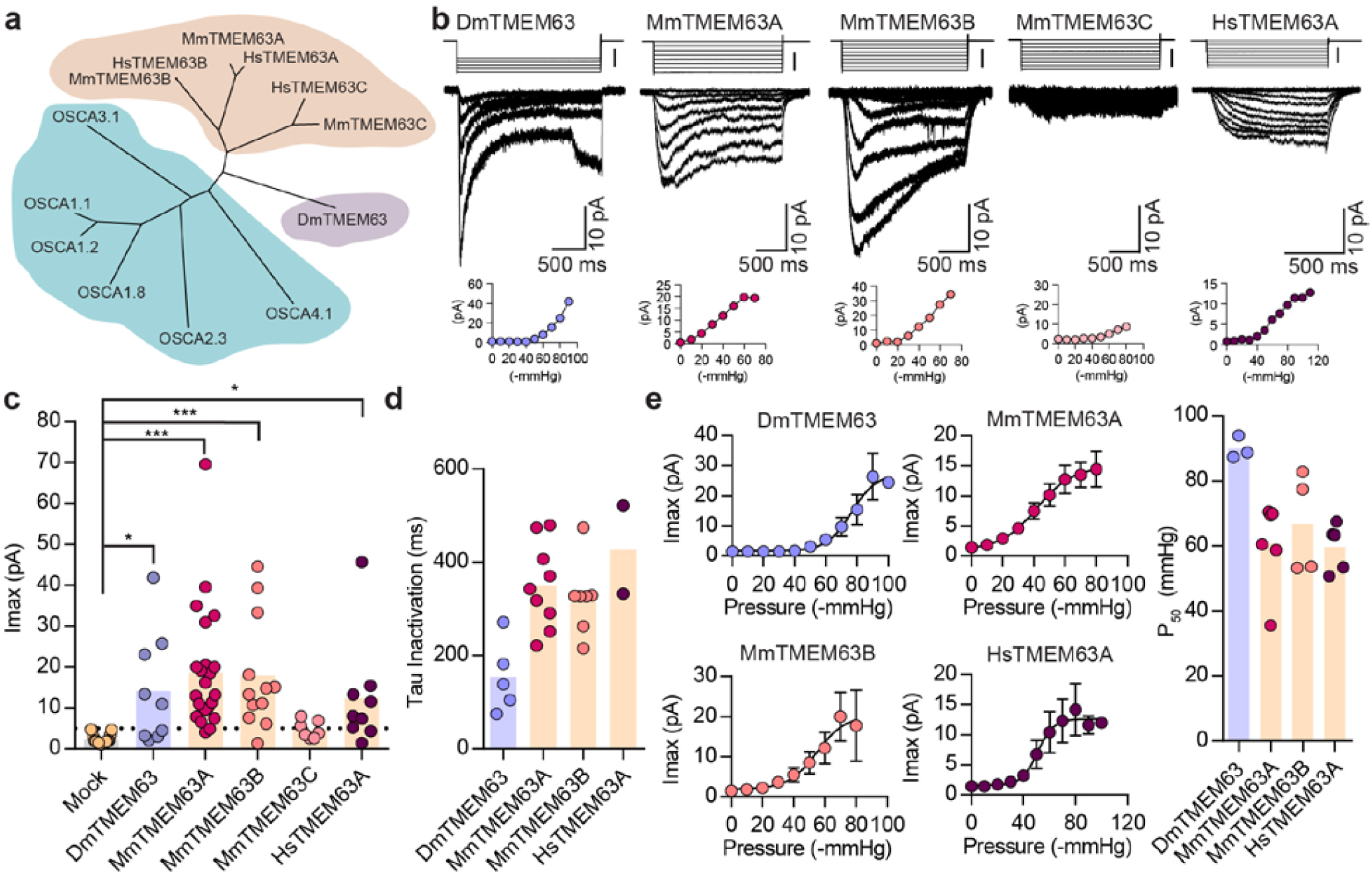
OSCA orthologues in flies and in mammals induce MA currents in HEKP1-KO cells. **a,** Phylogenetic tree illustrating the relationship between OSCA proteins across *Arabidopsis thaliana* (teal clade), *Drosophila melanogaster* (purple clade), and *Mus musculus* and *Homo sapiens* (orange clade). Sequences were aligned in MegAlign Pro and tree was generated in DrawTree. **b,** Representative stretch-activated currents induced by negative pipette pressure from cells expressing the indicated isoform. Corresponding pressure stimulus is illustrated above the current trace. Vertical scale bar: −50 mmHg. Pressure-response curve for the representative cell is illustrated below the trace. **c,** Maximal stretch-activated currents recorded from cells expressing mock plasmid (N=17) or the indicated TMEM63 isoform (DmTMEM63 (N=7), MmTMEM63A (N=23), MmTMEM63B (N=12), MmTMEM63C (N=7), and HsTMEM63A (N=6) (DmTMEM63: \**P*=0.023, HsTMEM63A: *P=0.015, \*\*\**P<*0.0001, Dunn’s multiple comparison test relative to mock plasmid). **d,** Inactivation time constant (ms) across individual cells from each isoform. Owing to the non-inactivating nature of HsTMEM63A currents, only 2 out of 9 cells could be reliably fit with an exponential curve (DmTMEM63 (N=5), MmTMEM63A (N=9), MmTMEM63B (N=7), and HsTMEM63A (N=2)). **e,** Left, average pressure-response curves fit with Boltzmann equation for the indicated isoform. Right, P_50_ values for individual cells across the four different TMEM63 isoforms (DmTMEM63 (N=3), MmTMEM63A (N=6), MmTMEM63B (N=4), and HsTMEM63A (N=5)).

P_50_ values for the TMEM63-induced stretch-activated currents were in the range of 60-80 mmHg, suggesting that like OSCAs, these proteins elicit high-threshold MA currents in HEK-P1KO cells (**Fig. 4e**). The stretch-activated single-channel currents induced by either DmTMEM63 or the mammalian TMEM63s are in the sub-picoampere range and were unresolvable, similar to OSCA2.3 (**Fig. 4b** vs. **Fig. 2b**). However, compared to plant OSCAs, MA currents induced by the TMEM63 family members had unique gating properties with several fold slower activation and inactivation kinetics (**Fig. 4d**, **Extended Data Table. 1**). MmTMEM63A stretch-activated currents are non-selective cationic, and gadolinium inhibits MmTMEM63A-induced MA currents by 75% (**Extended Data Fig. 6**). These results demonstrate that orthologues of OSCAs also induce stretch-activated currents when overexpressed in naïve cells, and that mechanosensitivity is conserved across the various isoforms in the OSCA/TMEM63 family.

We provide evidence that members of OSCA family are bona-fide, pore-forming mechanosensitive ion channels: (1) transfection of various members of the family from plants, flies, and mammals give rise to robust MA currents. Importantly, single-channel conductance of individual OSCAs are quite distinct, arguing that the OSCAs contribute to pore properties of these currents. (2) We directly show *in vitro* (proteoliposomes) that OSCA1.2 is an inherently mechanosensitive ion channel in the absence of other proteins. (3) The accompanying paper describes the high-resolution structure of OSCA1.2^12^, and demonstrates that this protein has similar architecture to the TMEM16 family of ion channels^26^. Furthermore, structure-guided mutagenesis verifies that a residue within the putative pore-forming region contributes to single-channel conductance. Therefore, with 15 different members present just in *Arabidopsis thaliana* (5/6 members we tested were mechanosensitive), OSCA/TMEM63 proteins potentially represent the largest family of mechanosensitive ion channels known to date. Although other eukaryotic ion channels have been previously described, none are conserved from plants to humans. PIEZOs are also present in plants and unicellular organisms; however, evidence that they are mechanosensitive in these species is still lacking.

The existence of 15 members of OSCAs in *Arabidopsis thaliana* raises the possibility that there might be redundancy, and will make it a challenge to assign function to these proteins individually. However, Yuan et al reported that under hyperosmotic stress mutant OSCA1.1 plants had stunted leaf and root growth^11^. These phenotypes could indeed be a consequence of impaired mechanotransduction. Expression profile of mouse TMEM63 genes in public databases (BioGPS.org) suggests expression in tissues that experience mechanical forces such as kidney and stomach. Future studies will investigate the contribution of members of OSCA and TMEM63 families to mechanotransduction in various species.

## Acknowledgements

We thank Drs. Kei Saotome, Jorg Grandl, Viktor Lukacs, Jose Santos, and Michael Bandell, and members of the Patapoutian lab for helpful discussions. We acknowledge Allain Fransisco, Meaghan Loud, Adam Coombs, and Jayanthi Mathur for technical support.

## Author contribution

S.E.M. and A.P. designed experiments. S.E.M performed and analyzed all electrophysiology experiments. A.E.D. performed and analyzed some whole-cell patch clamp experiments. T.W. and S.A.R.M. performed the cloning and molecular biology. S.C., A.E.D., S.E.M., and A.P. did bioinformatics analysis. S.J. purified and reconstituted proteins for the liposome experiments. S.E.M. and A.P. wrote the manuscript with input from other authors. A.E.D. is funded by National institute of Health (R21DE025329) and S.J. was supported by a Ray Thomas Edwards Foundation grant to A.B.W. This work was supported by National Institute of Health (NINDS) grant (R35NS105067) to A.P., who is also an investigator of the Howard Hughes Medical Institute.

## Methods

### Generation of clones

The different OSCA clones were gene synthesized (human codon optimized) from Genewiz. Sequences for *Arabidopsis thaliana* OSCA1.1 (At4g04340), OSCA1.2 (At4g22120), OSCA1.8 (At1g32090), OSCA2.3 (At3g01100), OSCA 3.1 (At1g30360), and OSCA4.1 (At4g35870) were downloaded from TAIR (www.arabiopsis.org). The synthesized cDNA was cloned into pIRES2-mCherry vector. In addition, OSCA1.1, OSCA1.2, and OSCA3.1 cDNA were cloned from *Arabidopsis thaliana* into pIRES2-mCherry vector. The OSCA1.1, OSCA1.2, and OSCA3.1 codon region were amplified from Arabidopsis cDNA with following primers:

OSCA1.1 Forward primer: ccgctagcgctaccggactcagatcATGGCAACACTTAAAGAC OSCA1.1 Reverse primer: gggcccgcggtaccgtcgactgcagCTAGACTTCTTTACCGTTAATAAC OSCA1.2 Forward primer: ccgctagcgctaccggactcagatcATGGCGACACTTCAGGATATTG OSCA1.2 Reverse primer: gggcccgcggtaccgtcgactgcagTTAGACTAGTTTACCACTAAAGGG OSCA3.1 Forward primer: gattaacagaagcttcccggCATGGAGTTTGGATCTTTTCTTGTG OSCA3.1 Reverse primer: gcccttgctcaccatgagctCAACGCCTGCTATTGCGTTG

pIRES2-mCherry plasmid was cut with FastDigest EcoRI and XhoI restriction enzyme (Thermo Fischer Scientific). Then the purified PCR product ligated into the digested plasmid using Gibson assembly kit (NEB). The sequence of genes verified by sequencing. The protein sequence downloaded from TAIR matched the sequence obtained from the plant. Furthermore, MA currents recorded from either clones were indistinguishable in their properties. Therefore, data for OSCA1.1, OSCA1.2, and OSCA3.1 was combined from gene synthesized cDNA and the cDNA sub-cloned from plants. *Drosophila melanogaster* TMEM63 (Dmel_CG11210, GenBank: AAF59136.1) was gene synthesized according to the sequence from GenBank. Mammalian TMEM63 clones were purchased from ORIGENE; MmTMEM63A (Cat No.: MR210748), MmTMEM63B (Cat No.: MR221527), MmTMEM63C (Cat No.: MC221684), and HsTMEM63A (Cat No.: RC206992). Some clones had a Myc tag when purchased, which was removed using QuickChange II XL site-directed mutagenesis kit according to the manufacturer’s instruction. The ORF of MmTMEM63A and MmTMEM63B was sub-cloned into pIRES2-mCherry vector, and MmTMEM63C was sub-cloned into pcDNA3.1-IRES-GFP. All clones were full-length sequence verified before testing.

### Cell culture and transient transfection

PIEZO1-knockout Human Embryonic Kidney 293T (HEK-P1KO) were used for all heterologous expression experiments. HEK-P1KO cells were generated using CRISPR–Cas9 nuclease genome editing technique as described previously^14^, and were negative for mycoplasma contamination. Cells were grown in Dulbecco’s Modified Eagle Medium (DMEM) containing 4.5 mg.ml^-1^ glucose, 10% fetal bovine serum, 50 units.ml^-1^ penicillin and 50 µg.ml^-1^ streptomycin. Cells were plated onto 12-mm round glass poly-D-lysine coated coverslips placed in 24-well plates and transfected using lipofectamine 2000 (Invitrogen) according to the manufacturer’s instruction. All plasmids were transfected at a concentration of 600 ng.ml^-1^. Cell were recorded from 24 to 48 hours after transfection. Since HsTMEM63A was in a non-fluorescent tagged vector, it was co-transfected with IRES-GFP or pIRES2-mCherry vector.

### Electrophysiology

Patch-clamp experiments in cells and in liposomes were performed in standard whole-cell, cell-attached, or excised patch (outside-out for cells, inside-out for liposomes) mode using Axopatch 200B amplifier (Axon Instruments). Some whole-cell recordings were done using Axon Multiclamp700A. Currents were sampled at 20 kHz and filtered at 2 kHz or 10kHz. Leak currents before mechanical stimulations were subtracted off-line from the current traces. Voltages were not corrected for a liquid junction potential (LJP) except for ion selectivity experiments. LJP was calculated using Clampex 10.6 software. All experiments were done at room temperature.

### Solutions

For whole-cell patch clamp recordings, recording electrodes had a resistance of 2-3 MΩ when filled with internal solution composed of (in mM) 133 CsCl, 1 CaCl_2_, 1 MgCl_2_, 5 EGTA, 10 HEPES (pH 7.3 with CsOH), 4 MgATP and 0.4 Na_2_GTP. The extracellular solution, also used as ios-osmotic solution, was composed of (in mM) 133 NaCl, 3 KCl, 2.5 CaCl_2_, 1 MgCl_2_, 10 HEPES (pH 7.3 with NaOH) and 10 glucose, 300 mmol/kg. Hyperosmolarity solution composed of (mM) 133 NaCl, 3 KCl, 2.5 CaCl_2_, 1 MgCl_2_, 10 HEPES (pH 7.3 with NaOH) and 300 Sorbitol, 620 mmol.kg^-1^.

For cell-attached patch clamp recordings, external solution used to zero the membrane potential consisted of (in mM) 140 KCl, 1 MgCl_2_, 10 glucose and 10 HEPES (pH 7.3 with KOH). Recording pipettes were of 1-3 MΩ resistance when filled with standard solution composed of (in mM) 130 mM NaCl, 5 KCl, 1 CaCl_2_, 1 MgCl_2_, 10 TEA-Cl and 10 HEPES (pH 7.3 with NaOH). For gadolinium inhibition experiments, 100 mM GdCl_3_ stock solution of was diluted in cell-attached pipette solution at a working concentration of 60 µM.

Ion selectivity experiments for OSCAs were performed in outside-out patch configurations. P_Cl_/P_Na_ was measured in extracellular solution composed of (in mM) 30 NaCl, 10 HEPES and 225 Sucrose (pH 7.3 with NaOH) and intracellular solution consisted of (in mM) 150 NaCl and 10 HEPES (pH 7.3 with NaOH). Ion selectivity experiments (**Extended Data Fig. 7**) on MmTMEM63A were performed in cell-attached patch clamp configuration. Independent cells were recorded under different conditions. NMDG-Cl solution consisted of (in mM) 150 NMDG,10 HEPES (pH 7.5). KCl solution consisted of (in mM) 150 KCl, 10 HEPES (pH 7.5). Cs-Meth solution consisted of (in mM): 149 Cs-methanesulphonate, 1 CsCl, 10 HEPES (pH 7.5).

### Mechanical stimulation

For whole-cell recordings, mechanical stimulation was achieved using a fire-polished glass pipette (tip diameter 3-4 µm) positioned at an angle of 80 º relative to the cell being recorded. Downward displacement of the probe towards the cell was driven by Clampex-controlled piezo-electric crystal microstage (E625 LVPZT Controller/Amplifier; Physik Instrumente). The probe had a velocity of 1 µm.ms^-1^ during the ramp phase of the command for forward movement and the stimulus was applied for 150 ms. To assess the mechanical sensitivity of a cell, a series of mechanical steps in 0.5 or 1 μm increments was applied every 10–20 s. Threshold was calculated as the differential (y-x) of the probe distance that first touches the cell (x) and the probe distance that induces the first channel response (y).

Macroscopic stretch-activated currents were recorded in the cell-attached or excised, outside-out patch clamp configuration. Membrane patches were stimulated with 500 ms (for OSCA clones) or 1 second or 2 second (for TMEM63 clones) negative or positive pressure pulses through the recording electrode using Clampex controlled pressure clamp HSPC-1 device (ALA-scientific), with inter-sweep duration of 1 min. Since TMEM63 had slower gating kinetics, longer stimulus duration was picked. Negative pressure was applied when patch was in cell-attached configuration, positive pressure was applied when patch was in the outside-out configuration. Activation time constant was determined by measuring 10-90% rise time (between baseline and peak) for currents at saturating pressure stimulus using Clampfit 10.6.

Stretch-activated single-channel currents were recorded in the cell-attached configuration. Since single-channel amplitude is independent of the pressure intensity, the most optimal pressure stimulation was used to elicit responses that allowed single-channel amplitude measurements. These stimulation values were largely dependent on the number of channels in a given patch of the recording cell. Single-channel amplitude at a given potential was measured from trace histograms of 5 to 10 repeated recordings. Histograms were fitted with Gaussian equations using Clampfit 10.6 software. Single-channel slope conductance for each individual cell was calculated from linear regression curve fit to single-channel I-V plots. Single-channel current for MscL in proteoliposomes was measured at one membrane voltage (30 mV) and conductance was calculated at that potential assuming 0 pA at 0 mV. In cells, OSCA1.2 single-channel amplitude in 200 mM KCl, 5 mM MOPS, pH 7.0 (KOH) was also measured at −80 mV (again assuming 0 pA at 0 mV) and used to calculate conductance.

### Hyperosmotic stimulation

Hyperosmolarity-induced currents were evoked by 2 different protocols^10,11^. 1) Once in the whole-cell-patch clamp configuration in iso-osmotic solution (300 mmol.kg^-1^) currents were recorded continuously in response to voltage ramps from −100 mV to +100 mV applied every 2-10s. Once a stable response was achieved, the bath solution was switched to hyperosmotic solution (620 mmol.kg^-1^) and currents were recorded for at least 5 additional minutes. Maximal response at −100 mV in iso-osmotic solution and hyper-osmotic solution were measured for each cell and plotted. 2) In the whole-cell patch clamp configuration whole-cell currents at - 80mV were recorded as cells were first exposed to 1 min of iso-osmotic (300 mmol.kg^-1^) solution followed by 5 mins of hyperosmotic solution (620 mmol.kg^-1^) and back to iso-osmotic solution. Maximal current response in the 3 conditions were plotted for each cell.

### Permeability ratio measurements

Reversal potential for each cell in the mentioned solution was determined by interpolation of the respective current-voltage data. Permeability ratios were calculated by using the following Goldman-Hodgkin-Katz (GHK) equations:

P_Cl_/P_Na_ ratios:

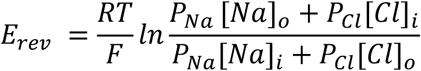

### OSCA1.2-PP-EGFP expression and purification

Codon optimized OSCA1.2 gene were cloned into vector pcDNA3.1. An EGFP tag was placed at the C terminus and connected to the gene via a PreScission Protease cleavable linker (LEVLFQGP) (OSCA1.2-PP-EGFP). A FLAG tag (DYKDDDDK) was added to the C terminus of EGFP with two intervening alanines as a linker. For protein expression and purification, four liters of HEK293F cells were grown in Freestyle 293 expression media to a density of 1.2-1.7×106 cells.mL^-1^. Each liter was transfected by combining 1mg.L^-1^ of the construct with 3 mg.L^-1^ of PEI in 30 mL of Opti-MEM and then adding the mix to the culture of cells. Transfected cells were grown for 48 hours and then pelleted, washed with ice cold PBS, flash frozen and stored at −80°C for future use. From this point forward, every step of the purification was carried out at 4°C unless otherwise stated. Pellets were thawed on ice, resuspended in 200 mL of solubilization buffer (25mM tris pH 8.0, 150mM NaCl, 1% Lauryl Maltose Neopentyl Glycol (LMNG), 0.1% cholesteryl hemisuccinate (CHS), 2µg.mL^-1^ leupeptin, 2µg.mL^-1^ aprotinin, 1mM PMSF, 2µM pepstatin, 2mM DTT) and stirred vigorously for 2-3 hours. Subsequently, insoluble material was pelleted via ultracentrifugation for 45 minutes at 30,000 rpm in a Type 70 Ti rotor. Batch binding of the supernatant was performed for 1 hour with 2mL of FLAG M2 affinity resin (Sigma) previously washed first with 0.1M glycine pH 3.5 and then with wash buffer (25 mM tris pH=8.0, 150mM NaCl, 0.01% LMNG, 0.001% CHS, 2mM DTT). Resin was placed in a gravity flow column and washed with 10CV of wash buffer. Protein was eluted using 2.5mL of elution buffer (wash buffer and 200 ug.mL^-1^ 3x FLAG peptide). Sample was concentrated using a 100kDa MWCO Amicon Ultra centrifugal filter. Concentrated protein was injected onto Shimadzu HPLC and size exclusion chromatography was performed using a Superose 6 Increase column and wash buffer. Fractions corresponding to OSCA1.2-PP-EGFP were concentrated to 2mg.mL^-1^.

### EcMscL expression and purification

EcMscL was purchased from Addgene (plasmid # 92418). EcMscL purification was done as previously described^27^, with the exception that we performed batch binding with Ni-NTA Agarose (QIAGEN).

### Proteoliposome reconstitution

Protocol for proteoliposome reconstitution was modified from previous publication^4^. Soybean polar lipid extract ((Avanti #541602) was completely desiccated and then resuspended in 200mM KCl, 5mM MOPS, pH 7.0 for a final concentration of 10mg.mL^-1^. The mixture was then bath sonicated for 3 cycles of 2 minutes sonication followed by 2 minutes wait. Liposomes were aliquoted and frozen for future use. Liposomes were thawed and supplemented with DDM for a final concentration of 1.5 mM DDM. Protein was diluted to 2mg.mL^-1^ and was added to liposomes in a 1:100 ratio (protein:lipid). Mixture was placed on ice for 5 minutes and then incubated at room temperature for 20 min with rotation. 10 mg of previously washed biobeads (one methanol wash, two water washes and one wash with 200 mM KCl, 5 mM MOPS, pH 7.0) were added to the mixture and incubated for 1 hour with rotation at room temperature. Biobeads were removed and a second set of 10 mg of biobeads was added and incubated for 30 minutes. Biobeads were removed and mixture was centrifuge at 60,000 rpm for 60 minutes at 8ºC in a Beckman Coulter Optima TLX ultracentrifuge.

### Proteoliposome recordings

Protocol used to record stretch-activated currents from proteoliposomes was adapted from Coste et al., 2012^4^. The proteoliposome pellet was re-suspended in 40 μl of buffer containing 200 mM KCl, 5 mM MOPS (pH7.0) and used to place two 20 μl drops on a cover slide. The samples were dried under vacuum for >16 hours. Samples were then hydrated with 25 μl of the same buffer and allowed to sit for 2 hours at 4°C before starting recordings. 2-3 μl of proteoliposomes were withdrawn from the edge of the spots on the cover slide and transferred to the recording chamber. After 5 min, the chamber was slowly filled with recording solution. Multi-GΩ seals were made to proteoliposomes immobilized at the bottom of the recording chamber. At that time, the proteoliposome patch was excised to create an inside-out patch. Pipette and bath solution contained (in mM) 200 KCl, 5 MOPS titrated to pH 7.0 with KOH.

### Data availability

Accession codes where applicable have been provided. All other data are available upon request to the corresponding author.

---

**Extended Data Figure 1:**
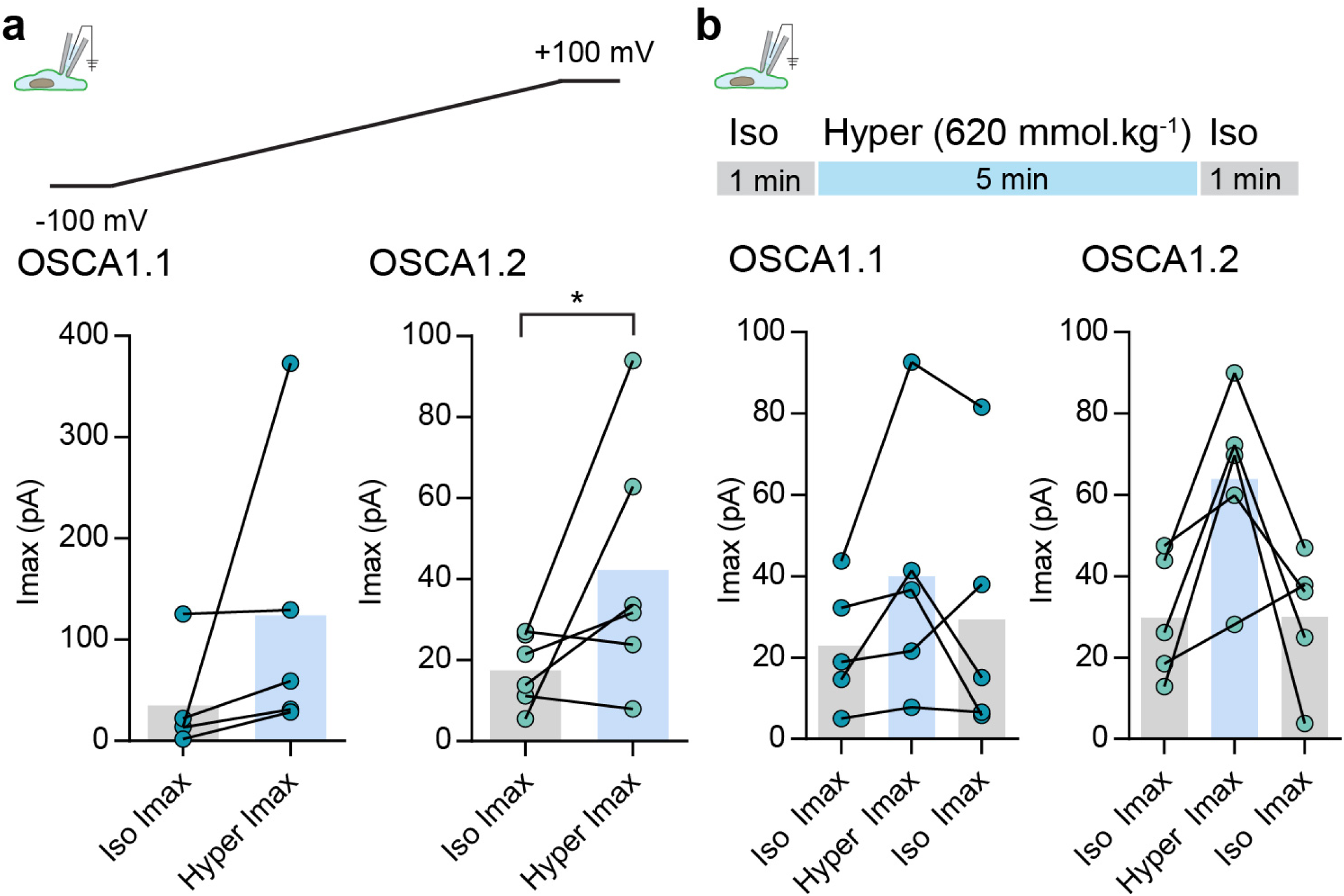
Hyperosmolarity-evoked currents from OSCA1.1- and OSCA1.2-transfected HEK-P1KO cells. **a,** Maximal current response recorded at −100 mV from cells expressing OSCA1.1 or OSCA1.2 in iso-osmotic (300 mmol.kg^-1^) or hyperosmotic solutions (620 mmol.kg^-1^). In the whole-cell patch clamp mode currents were recorded in response to voltage ramps between −100 mV and +100 mV. Maximal response from each cell in iso-and hyper-osmotic solution is represented as scatter plot, bars represent population mean. OSCA1.1: Iso = 35 ± 22 pA; Hyper = 124 ± 64 pA (N=6), OSCA1.2: Iso = 17 ± 3 pA; Hyper = 42 ± 10 pA (N=6) (\**P*=0.0317, Mann-Whitney test). **b**, Hyperosmotic currents were induced by applying a 5 minute stimulus of hypertonic solution (620 mmol.kg^-1^) solution. Currents were continuously recorded at - 80mV. Maximal current for individual cells in the indicated condition is represented as scatter plot, bars represent population mean. OSCA1.1: Iso = 23 ± 6.8 pA; Hyper = 40 ± 14 pA; Iso = 30 ± 14 pA (N=5), OSCA1.2: Iso = 30 ± 6.8 pA; Hyper = 64 ± 10 pA; Iso = 30 ± 7 pA (N=5).

**Extended Data Figure 2.**
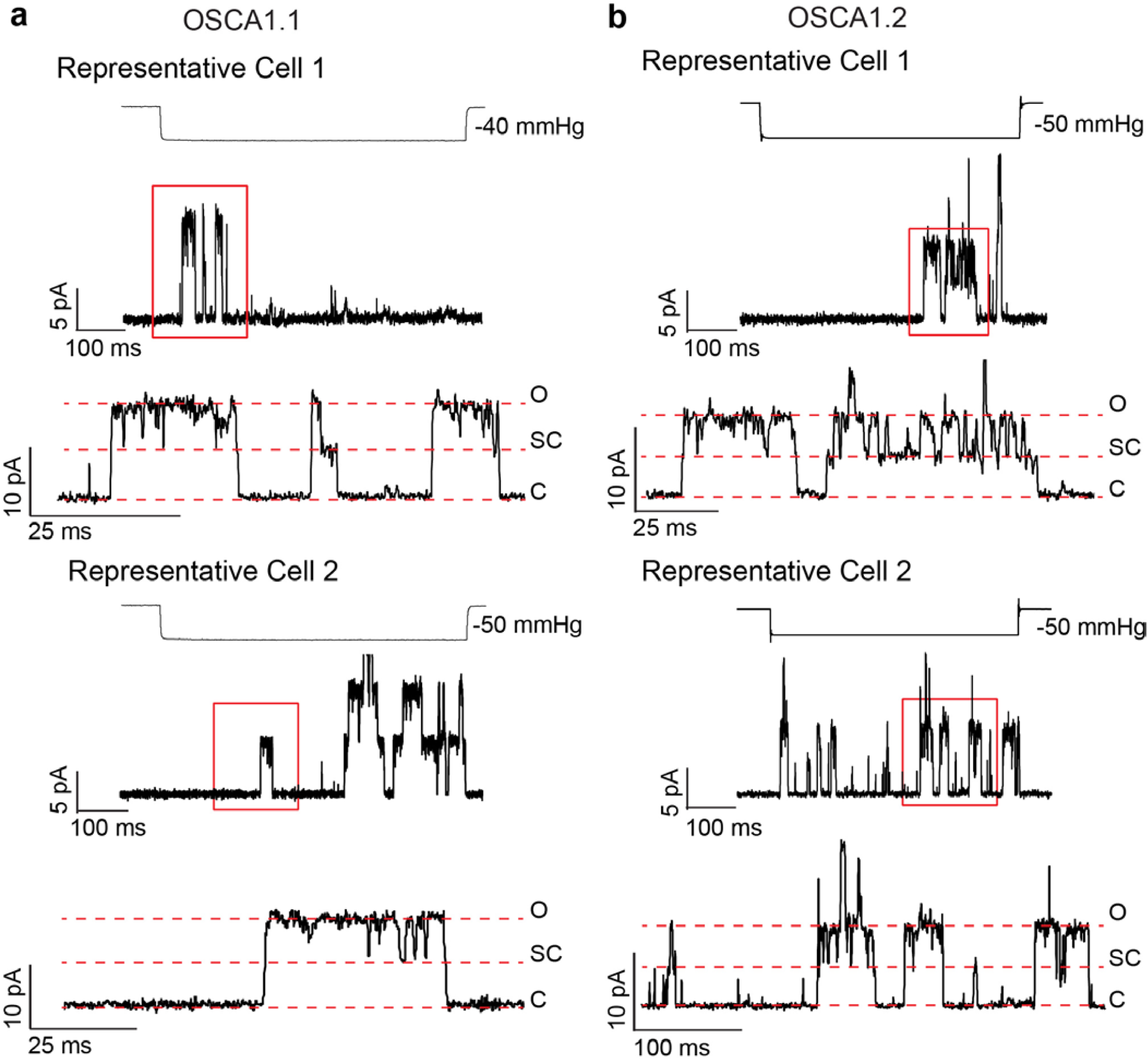
Subconductance states in OSCA1.1- and OSCA1.2-dependent stretch-activated single-channel traces. Representative traces from two different cells for OSCA1.1 (**a**) and OSCA1.2 (**b**). Channel openings are upward deflections, and currents were recorded at −80 mV but are illustrated as upward currents. For each representative cell, top trace illustrates a longer time scale and channel opening in response to stretch. The corresponding pressure-stimulus is illustrated above. The bottom trace highlights the subconductance state. Red box indicates the expanded region in the bottom trace. C: closed, SC: subconductance, O: open.

**Extended Data Figure 3.**
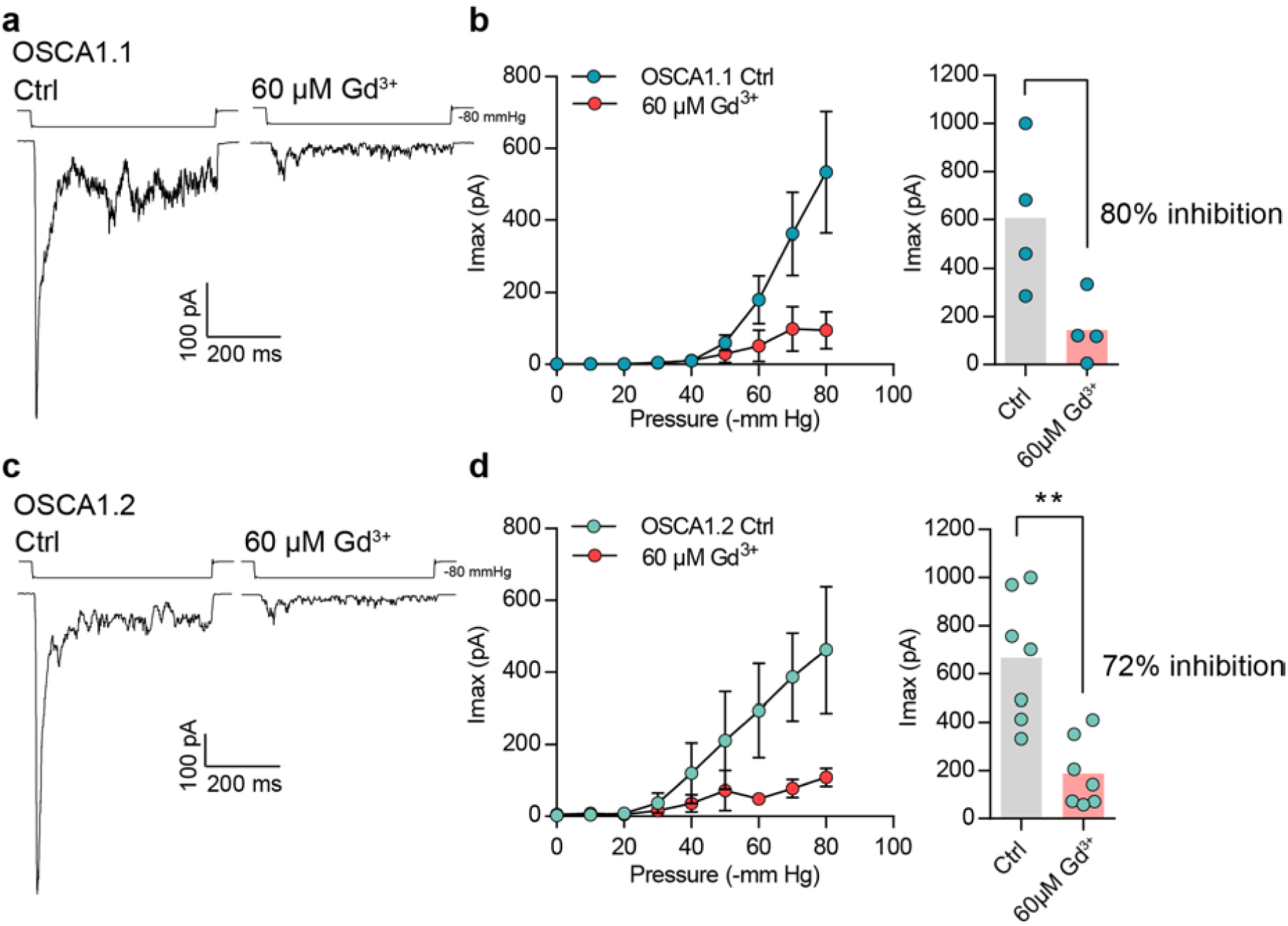
Gadolinium block of OSCA1.1- and OSCA1.2-dependent MA currents. **a, c,** Representative trace of stretch-activated currents at −80 mV in the cell-attached patch clamp mode. Currents were evoked by −80 mmHg of pressure in the presence of physiological pipette solution (Ctrl) or physiological pipette solution with 60 µM Gd^3+^ from cells expressing OSCA1.1 (**a**) or OSCA1.2 (**c**). Traces on left and right are from two independent cells. **b**, **d**, Left, average pressure-response curves of stretch-activated currents recorded from OSCA1.1-(N=4) (**b**) or OSCA1.2-(N=7) (**d**) expressing cells in Ctrl solution or Ctrl+60 µM Gd^3+^. Right, Average maximal current across individual cells without (OSCA1.1: Imax= 606 ± 154 pA (N=4); OSCA1.2: Imax= 666 ± 100 pA (N=7)) or with Gd^3+^ (OSCA1.1: Imax= 144 ± 68 pA (N=4); OSCA1.2: Imax= 187 ± 54 pA (N=7)). Bars represent population mean. Gd^3+^ blocks OSCA1.1 and OSCA1.2 maximal current responses by 80% and 72%, respectively (\*\**P*=0.0023, Mann-Whitney test).

**Extended Data Figure 4:**
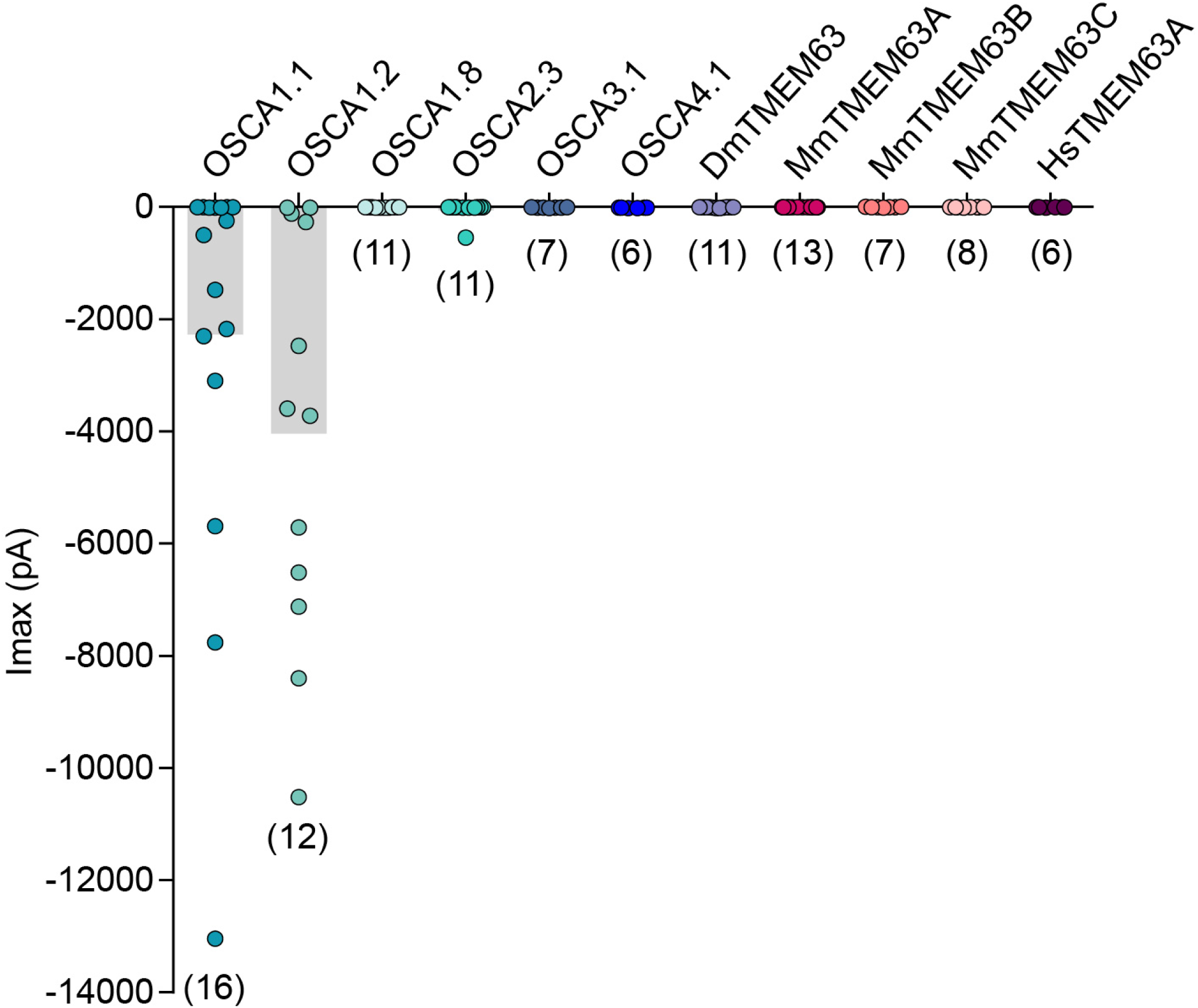
Membrane indentation-induced MA currents in HEK-P1KO cells transfected with OSCA and TMEM63 isoforms. MA whole-cell maximal inward currents from individual cells expressing the indicated isoform. The last current response before losing the cell is reported. Currents were elicited at - 80mV membrane potential. Numbers in parenthesis indicate number of cells tested per isoform. Bars represent population mean.

**Extended Data Figure 5.**
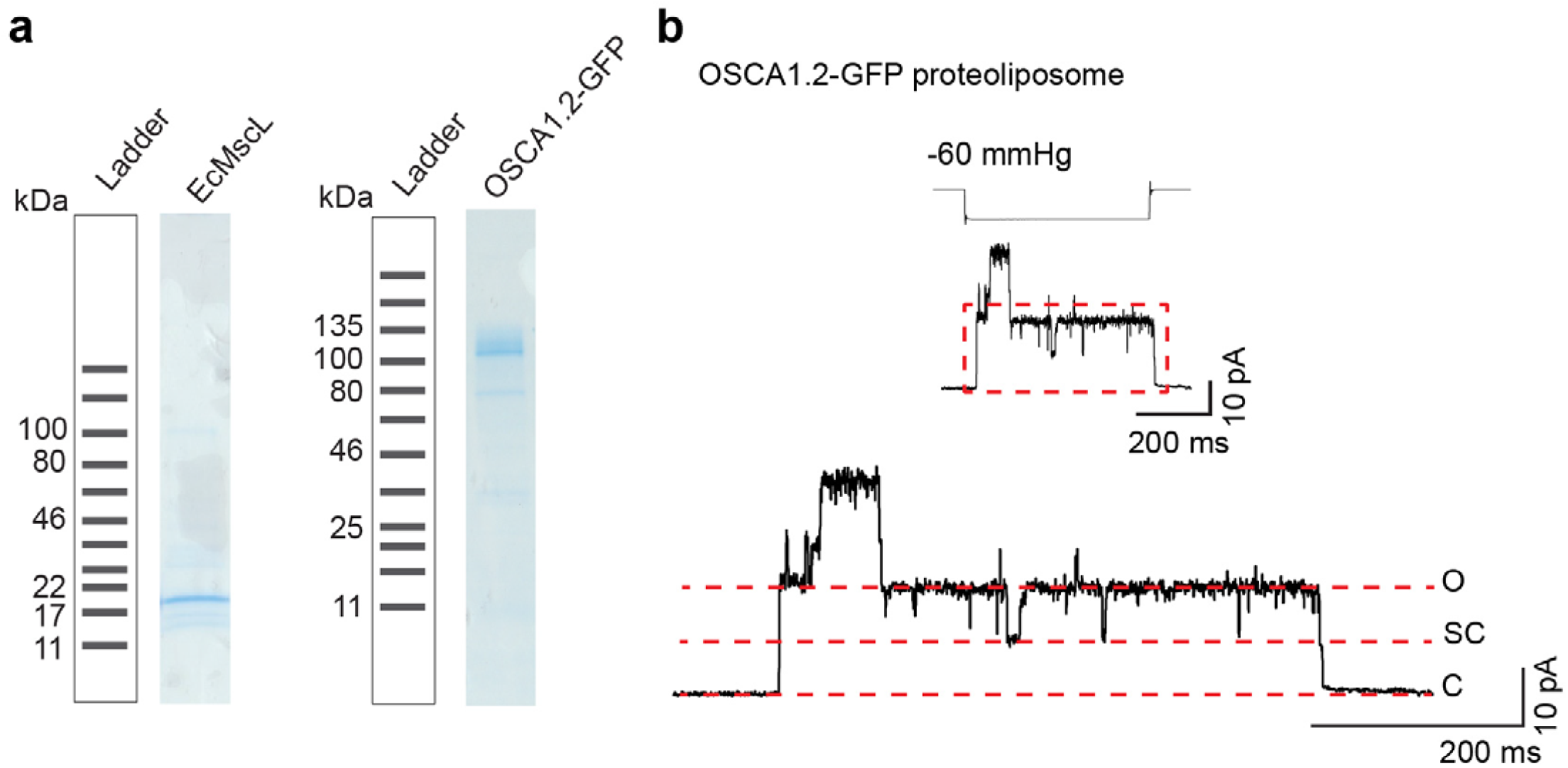
Purified OSCA1.2 induces stretch-activated subconductance states when reconstituted in liposomes. **a,** SDS-PAGE showing purification of, EcMscL and OSCA1.2. Samples are taken after eluting protein from resin and before concentrating it for further steps (proteoliposome reconstitution for EcMscL or after size exclusion chromatography for OSCA1.2). Expected monomer size: EcMscL 17 kDa and OSCA1.2-GFP 117 kDa. **b**, Top, representative stretch-activated single-channel currents (−80 mV) from OSCA1.2-GFP reconstituted in liposomes. Stimulus trace is depicted above the current trace. Bottom, enlarged current trace to illustrate sub-conductance states. Channel openings are upward deflections. C: closed, SC: sub-conductance, O: open.

**Extended Data Figure 6.**
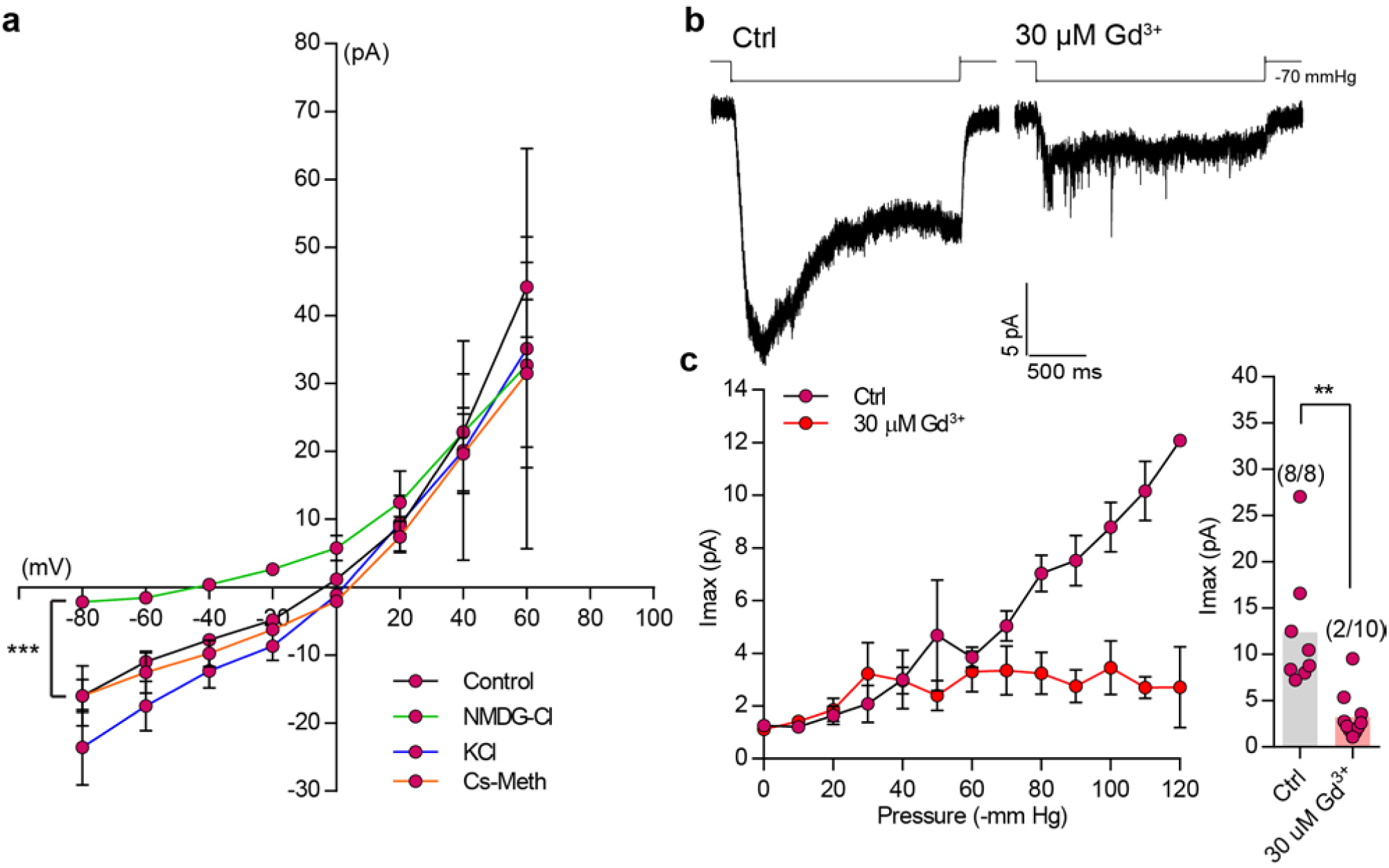
Ion selectivity and gadolinium block of MmTMEM63A-induced MA currents. **a,** I-V relation curves for stretch-activated currents recorded in the cell-attached patch clamp mode from cells expressing MmTMEM63A in the in indicated ion solution (Control: N=11, NMDG-Cl: N=10, KCl: N=4, and Cs-Meth: N=7, *** *P*<0.0001, Two-way ANOVA, not RM between control and NMDG-Cl). **b,** Representative trace of stretch-activated current evoked by −70 mmHg of pressure in the presence of physiological pipette solution (Ctrl) or physiological pipette solution with 30 µM Gd^3+^. Traces on left and right are from two independent cells. **c**, Left, average pressure-response curves of stretch-activated currents recorded from MmTMEM63A expressing cells in Ctrl solution (N=8) or Ctrl+30 µM Gd^3+^ (N=10). Right, Average maximal current across individual cells with or without Gd^3+^. Bars represent population mean. Numbers in parenthesis denote cells with a response/total number of cells tested.

**Extended Data Table 1.** Biophysical properties of OSCA and TMEM63 isoforms.

| Isoforms | Whole-cell/Poke |  |  | Cell-attached/Stretch |  |  |  |  |
| --- | --- | --- | --- | --- | --- | --- | --- | --- |
|  | I<sub>max</sub> (pA) | Inactivation tau (ms) | Threshold (μm) | I<sub>max</sub> (pA) | Activation tau (ms) | Inactivation tau (ms) | P<sub>50</sub> (mmHg) | Channel conductance (pS) |
| Mock | 4.2 ± 0.5 (10) | - | - | 2.32 ± 0.3 (17) | - | - | - | - |
| Mm PIEZO1 | 3045 ± 875 (5) | 18.7 ± 1.7 (5) | 4.0 ± 0.6 (5) | 122 ± 40 (5) | 9.0 ± 1.0 (5) | 28.8 ± 2.0 (5) | 24.0 ± 3.6 (4) | - |
| OSCA1.1 | 2271 ± 918 (16) | 10.0 ± 1.3 (8) | 8.6 ± 0.9 (9) | 563 ± 68 (11) | 6.7 ± 1.0 (11) | 24.0 ± 3.0 (8) | 58.5 ± 2.4 (6) | 184.4 ± 4.4 (5) |
| OSCA1.2 | 4039 ± 1046 (12) | 10.4 ± 1.7 (9) | 6.3 ± 0.7 (10) | 576 ± 65 (14) | 5.5 ± 0.5 (14) | 24.6 ± 4.7 (9) | 54.5 ± 2.2 (10) | 121.8 ± 3.4 (4) |
| OSCA1.8 | 7.1 ± 0.7 (11) |  |  | 347 ± 56 (7) | 14.0 ± 2.4 (7) | 67.0 ± 13.0 (6) | 79.3 ± 9.9 (6) | 46.6 ± 2.8 (4) |
| OSCA2.3 | 54.34 ± 48.9 (11) |  |  | 31.3 ± 11 (7) | 18.8 ± 3.2 (6) | 110.0 ± 70.0 (6) | 59.7 ± 3.8 (7) | n.d. |
| OSCA3.1 | 9.7 ± 1.4 (7) |  |  | 306 ± 47 (12) | 5.6 ± 1.4 (8) | 18.5 ± 2.5 (11) | 44.5 ± 3.2 (5) | 24.9 ± 3.4 (4) |
| OSCA4.1 | 13.3 ± 2.0 (6) |  |  | 1.2 ± 0.05 (6) | - | - | - | - |
| Dm TMEM63 | 9.8 ± 2.0 (11) |  |  | 14.2 ± 4.5 (9) | 22.6 ± 4.3 (5) | 154 ± 34 (5) | 90 ± 2 (3) | n.d. |
| Mm TMEM63A | 2.0 ± 1.2 (13) |  |  | 18.75 ± 3.0 (23) | 126 ± 19 (8) | 323 ± 30 (9) | 60 ± 5.4 (6) | n.d. |
| Mm TMEM63B | 3.8 ± 1.8 (7) |  |  | 17.97 ± 3.9 (12) | 245 ± 38 (7) | 323 ± 30 (7) | 66 ± 7.8 (4) | n.d. |
| Mm TMEM63C | 4.5 ± 0.9 (8) |  |  | 4.73 ± 0.8 (7) | - | - | - | - |
| Hs TMEM63A | 4.2 ± 0.7 (6) |  |  | 12.4 ± 4.4 (9) | 188 ± 18 (5) | 427 ± 94 (2) | 59.7 ± 3.2 (5) | n.d. |

**Note:** n.d.: not determined, Ns are indicated in parenthesis. I<sub>max</sub> value is reported for the last indentation or stretch stimulus before losing the cell. Activation is reported as 10-90% rise of stretch-activated current at saturating stimulus. Inactivation time constant for stretch-activated currents are reported in the range of -60 to -80 mmHg stimulus-pressure.

**Extended Data Table 2.** Percent identity in protein sequence among OSCA and TMEM63 isoforms.

| Isoforms | OSCA 1.1 | OSCA 1.2 | OSCA 1.8 | OSCA 2.3 | OSCA 3.1 | OSCA 4.1 | Dm TMEM63 | Mm TMEM63A | Mm TMEM63B | Mm TMEM63C | Hs TMEM63A |
| --- | --- | --- | --- | --- | --- | --- | --- | --- | --- | --- | --- |
| OSCA 1.1 | 100 | 84.81 | 58 | 32.76 | 29.09 | 18.82 | 18.07 | 19.19 | 19.32 | 17.55 | 20.11 |
| OSCA 1.2 | - | 100 | 58.39 | 34.47 | 28.95 | 19.42 | 18.40 | 20.09 | 19.52 | 17.75 | 20.43 |
| OSCA 1.8 | - | - | 100 | 32.24 | 31.79 | 18.00 | 18.32 | 19.26 | 19.06 | 19.15 | 19.46 |
| OSCA 2.3 | - | - | - | 100 | 23.80 | 17.30 | 17.012 | 19.57 | 18.17 | 17.88 | 19.41 |
| OSCA 3.1 | - | - | - | - | 100 | 18.39 | 18.92 | 22.36 | 20.33 | 19.79 | 21.75 |
| OSCA 4.1 | - | - | - | - | - | 100 | 18.97 | 21.27 | 21.30 | 19.68 | 21.42 |
| Dm TMEM63 | - | - | - | - | - | - | 100 | 32.58 | 32.79 | 30.78 | 32.59 |
| Mm TMEM63A | - | - | - | - | - | - | - | 100 | 57.16 | 41.28 | 90.05 |
| Mm TMEM63B | - | - | - | - | - | - | - | - | 100 | 44.46 | 57.58 |
| Mm TMEM63C | - | - | - | - | - | - | - | - | - | 100 | 40.73 |
| Hs TMEM63A | - | - | - | - | - | - | - | - | - | - | 100 |

## Reference

1. Ranade, Sanjeev S., Syeda, R. & Patapoutian, A. Mechanically Activated Ion Channels. Neuron 87, 1162–1179, doi:http://dx.doi.org/10.1016/j.neuron.2015.08.032 (2015).

2. Árnadóttir, J. & Chalfie, M. Eukaryotic Mechanosensitive Channels. Annual Review of Biophysics 39, 111–137, doi:doi:10.1146/annurev.biophys.37.032807.125836 (2010).

3. Coste, B. et al. Piezo1 and Piezo2 Are Essential Components of Distinct Mechanically Activated Cation Channels. Science 330, 55–60, doi:10.1126/science.1193270 (2010).

4. Coste, B. et al. Piezo proteins are pore-forming subunits of mechanically activated channels. Nature 483, 176, doi:10.1038/nature10812 https://www.nature.com/articles/nature10812#supplementary-information (2012).

5. Murthy, S. E., Dubin, A. E. & Patapoutian, A. Piezos thrive under pressure: mechanically activated ion channels in health and disease. Nature Reviews Molecular Cell Biology 18, 771, doi:10.1038/nrm.2017.92 (2017).

6. Hamant, O. Widespread mechanosensing controls the structure behind the architecture in plants. Current Opinion in Plant Biology 16, 654–660, doi:https://doi.org/10.1016/j.pbi.2013.06.006 (2013).

7. Hamant, O. & Haswell, E. S. Life behind the wall: sensing mechanical cues in plants. BMC Biology 15, 59, doi:10.1186/s12915-017-0403-5 (2017).

8. Hamilton, E. S. et al. Mechanosensitive channel MSL8 regulates osmotic forces during pollen hydration and germination. Science 350, 438–441, doi:10.1126/science.aac6014 (2015).

9. Cox, C. D., Nakayama, Y., Nomura, T. & Martinac, B. The evolutionary ‘tinkering’ of MscS-like channels: generation of structural and functional diversity. Pflügers Archiv - European Journal of Physiology 467, 3–13, doi:10.1007/s00424-014-1522-2 (2015).

10. Hou, C. et al. DUF221 proteins are a family of osmosensitive calcium-permeable cation channels conserved across eukaryotes. Cell Research 24, 632, doi:10.1038/cr.2014.14 https://www.nature.com/articles/cr201414#supplementary-information (2014).

11. Yuan, F. et al. OSCA1 mediates osmotic-stress-evoked Ca2+ increases vital for osmosensing in Arabidopsis. Nature 514, 367, doi:10.1038/nature13593 https://www.nature.com/articles/nature13593#supplementary-information (2014).

12. Sebastian Jojoa-Cruz1, K. S., Swetha E. Murthy, Che Chun (Alex) Tsui, Mark S. P. Sansom, Ardem Patapoutian, Andrew B. Ward. Cryo-EM structure of the mechanically activated ion channel OSCA1.2. Submitted (2018).

13. Dubin, A. E. et al. Endogenous Piezo1 Can Confound Mechanically Activated Channel Identification and Characterization. Neuron 94, 266–270.e263, doi:https://doi.org/10.1016/j.neuron.2017.03.039 (2017).

14. Lukacs, V. et al. Impaired PIEZO1 function in patients with a novel autosomal recessive congenital lymphatic dysplasia. Nature Communications 6, 8329, doi:10.1038/ncomms9329 (2015).

15. Sachs, F. Stretch-Activated Ion Channels: What Are They? Physiology (Bethesda, Md.) 25, 50–56, doi:10.1152/physiol.00042.2009 (2010).

16. Coste, B. et al. Piezo1 ion channel pore properties are dictated by C-terminal region. Nat Commun 6, doi:10.1038/ncomms8223 (2015).

17. Dani, J. A. & Fox, J. A. Examination of subconductance levels arising from a single ion channel. Journal of Theoretical Biology 153, 401–423, doi:https://doi.org/10.1016/S0022-5193(05)80578-8 (1991).

18. Hartzell, H. C. & Whitlock, J. M. TMEM16 chloride channels are two-faced. The Journal of General Physiology 148, 367–373, doi:10.1085/jgp.201611686 (2016).

19. Miller, C. Open-state substructure of single chloride channels from Torpedo electroplax. Philosophical Transactions of the Royal Society of London. B, Biological Sciences 299, 401–411, doi:10.1098/rstb.1982.0140 (1982).

20. Häse, C. C., Le Dain, A. C. & Martinac, B. Purification and Functional Reconstitution of the Recombinant Large Mechanosensitive Ion Channel (MscL) of Escherichia coli. Journal of Biological Chemistry 270, 18329–18334, doi:10.1074/jbc.270.31.18329 (1995).

21. Kloda, A. & Martinac, B. Common evolutionary origins of mechanosensitive ion channels in Archaea, Bacteria and cell-walled Eukarya. Archaea 1, 35–44 (2002).

22. Brohawn, S. G., Su, Z. & MacKinnon, R. Mechanosensitivity is mediated directly by the lipid membrane in TRAAK and TREK1 K^+^ channels. Proceedings of the National Academy of Sciences 111, 3614–3619, doi:10.1073/pnas.1320768111 (2014).

23. Syeda, R. et al. Piezo1 Channels Are Inherently Mechanosensitive. Cell Reports 17, 1739–1746, doi:https://doi.org/10.1016/j.celrep.2016.10.033 (2016).

24. Anishkin, A., Loukin, S. H., Teng, J. & Kung, C. Feeling the hidden mechanical forces in lipid bilayer is an original sense. Proceedings of the National Academy of Sciences 111, 7898–7905, doi:10.1073/pnas.1313364111 (2014).

25. Xin, Z., Xiaojuan, Y., Yuanhu, L., Ping, Z. & Xin, N. Co-expression of mouse TMEM63A, TMEM63B and TMEM63C confers hyperosmolarity activated ion currents in HEK293 cells. Cell Biochemistry and Function 34, 238–241, doi:doi:10.1002/cbf.3185 (2016).

26. Whitlock, J. M. & Hartzell, H. C. Anoctamins/TMEM16 Proteins: Chloride Channels Flirting with Lipids and Extracellular Vesicles. Annual Review of Physiology 79, 119–143, doi:10.1146/annurev-physiol-022516-034031 (2017).

27. Rosholm, K. R. et al. Activation of the mechanosensitive ion channel MscL by mechanical stimulation of supported Droplet-Hydrogel bilayers. Scientific Reports 7, 45180, doi:10.1038/srep45180 https://www.nature.com/articles/srep45180#supplementary-information (2017).

